# From Sensory Details to Verbal Codes: Visual Memory Reinstatement of Location and Color Features

**DOI:** 10.64898/2026.01.27.701888

**Authors:** Xinhao Wang, Maxi Becker, Rasha Abdel Rahman, Simon W. Davis, Roberto Cabeza

## Abstract

Successful episodic memory depends on the reinstatement of encoding content and processes during retrieval. However, it remains unclear how such reinstatement (1) differs for various perceptual features, (2) supports subjective versus objective memory retrieval, and (3) relies on linguistic information. To investigate these three issues, we designed an fMRI study in which participants encoded and recalled the colors and spatial locations of object pictures, and provided subjective vividness and objective precision measures. We used encoding-retrieval similarity (ERS) to identify feature-level reinstatement for location or color features, and trial-specific reinstatement for individual location trials or individual color trials. The study yielded three main findings. First, feature-level reinstatement in frontoparietal and visual regions supported both location and color memory, whereas trial-specific reinstatement in early visual cortex contributed to location memory and feature-level reinstatement in lateral/anterior temporal cortices, to color memory. Second, trial-specific reinstatement in early visual cortex supported both location vividness and precision, while feature-level reinstatement in inferior frontal gyrus contributed to color vividness, and trial-specific reinstatement in posterior inferior temporal gyrus (including the color area), to color precision. Finally, color name richness (the number of names associated with a particular color) enhanced color memory precision and modulated trial-specific color reinstatement in right anterior inferior temporal gyrus. Together, these findings suggest that perceptual and linguistic reinstatement play complementary roles in visual episodic memory: perceptual replay provides fine-grained sensory details, whereas linguistic replay can scaffold and refine these representations, especially for features that are easier to verbalize such as colors.

**Significance Statement:** Neural reinstatement has been associated with the quality of episodic memory for complex visual images. However, it remains unclear how neural reinstatement varies across perceptual features, memory subjectivity, and linguistic accessibility. Here, we examine how neural reinstatement supports memory for color and location, whether it plays different roles in memory vividness versus precision, and the role of language in color memory. We showed that location memory relies primarily on trial-specific reinstatement in early visual cortex, supporting precise and vivid recall, whereas color memory is supported by a more complex and complementary interplay between perceptual and linguistic reinstatement. Together, these results advance our understanding of how distinct reinstatement mechanisms support the precision and vividness of episodic memory for basic visual features.

## Introduction

A popular hypothesis in cognitive memory research is that successful retrieval depends on the recapitulation of encoding processes and representations (Morris et al., 1977). Consistent with this hypothesis, functional neuroimaging studies have shown that brain regions activated at encoding are reactivated during retrieval, and this reactivation predicts memory success (Buckner & Wheeler, 2001; Rugg et al., 2008; Xue, 2018). In representational fMRI studies, neural reinstatement is typically quantified by the similarity of multivoxel activation patterns during encoding and retrieval, or *encoding-retrieval similarity* (ERS) (Ritchey et al., 2013; Wing et al., 2015). Multiple studies have shown that ERS in ventral temporal and medial temporal regions is associated with successful memory performance (Rau et al., 2025; Ritchey et al., 2013; Staresina et al., 2012; Wing et al., 2015), confirming the behavioral significance of neural reinstatement. Yet, very little is known on how reinstatement (1) differs for various perceptual features, (2) supports subjective versus objective memory measures, and (3) is affected by linguistic processes. The current study therefore had three goals.

### Our first goal was to compare neural reinstatement for distinct perceptual features, namely location vs. color episodic memory

Remembering basic visual features, such as location and color, supports rich, detailed episodic memory and guides everyday behavior. However, most fMRI studies on episodic reinstatement have emphasized complex visual scenes or objects (Lohnas et al., 2018; Rau et al., 2025; Wing et al., 2015), leaving simple visual features relatively understudied. Although fMRI studies on visual short-term memory have examined the reinstatement of location and color memory (Adam et al., 2022; Serences, 2016), studies on episodic memory recapitulation of these features are scarce (see however, Hou et al., 2025; Richter et al., 2016).

### Our second goal was to compare neural reinstatement supporting subjective vividness vs. objective precision

In episodic memory reactivation studies using scenes or objects, participants typically rate the subjective vividness of the remembered image (Hebscher et al., 2021; Qin et al., 2011; Wing et al., 2015). In contrast, in fMRI or EEG studies of short-term memory for simple visual features, such as color or location, participants often report the value of the encoded feature along a continuous scale, yielding an objective precision measure (Bays et al., 2009). In addition to linking these two different memory domains, comparing the neural reinstatement underlying vividness and precision is important, because subjective and objective retrieval may rely on distinct forms of reactivated information.

### Finally, our third goal was to examine how language shapes color memory reinstatement

Location and color differ in their reliance on language in everyday life, with color being a feature whose perception is known to be influenced by language (Lupyan et al., 2020). Behaviorally, some colors can be described using more verbal labels than others (Hasantash & Afraz, 2020), which we term here “color name richness.” One important difference between perceptual processing in human and non-human animals is that human’s perception is almost always affected by semantic and linguistic knowledge (Lupyan et al., 2020). Given the sensitivity of color processing to language, the reinstatement of color memory is an ideal test bed for examining the influence of language on memory processes in humans.

To address these goals, we collected fMRI while participants encoded either the color or the location of everyday objects (Figure 1) and then retrieved these features, while reporting vividness ratings and continuous feature values (precision). For *Goal 1*, we compared ERS for location versus color, and predicted that memory for location and color would be supported by ERS in dorsal vs. ventral visual stream, respectively (Ungerleider & Mishkin, 1982). For *Goal 2*, we compared ERS for vividness versus precision, and predicted that trial-specific ERS, which should reflect fine-grained feature variation, would more strongly track precision than vividness. For *Goal 3*, we conducted a normative study quantifying the number of distinct names for each hue (“color name richness”), and then tested how name richness modulated color reinstatement and memory performance.

**Figure 1.**
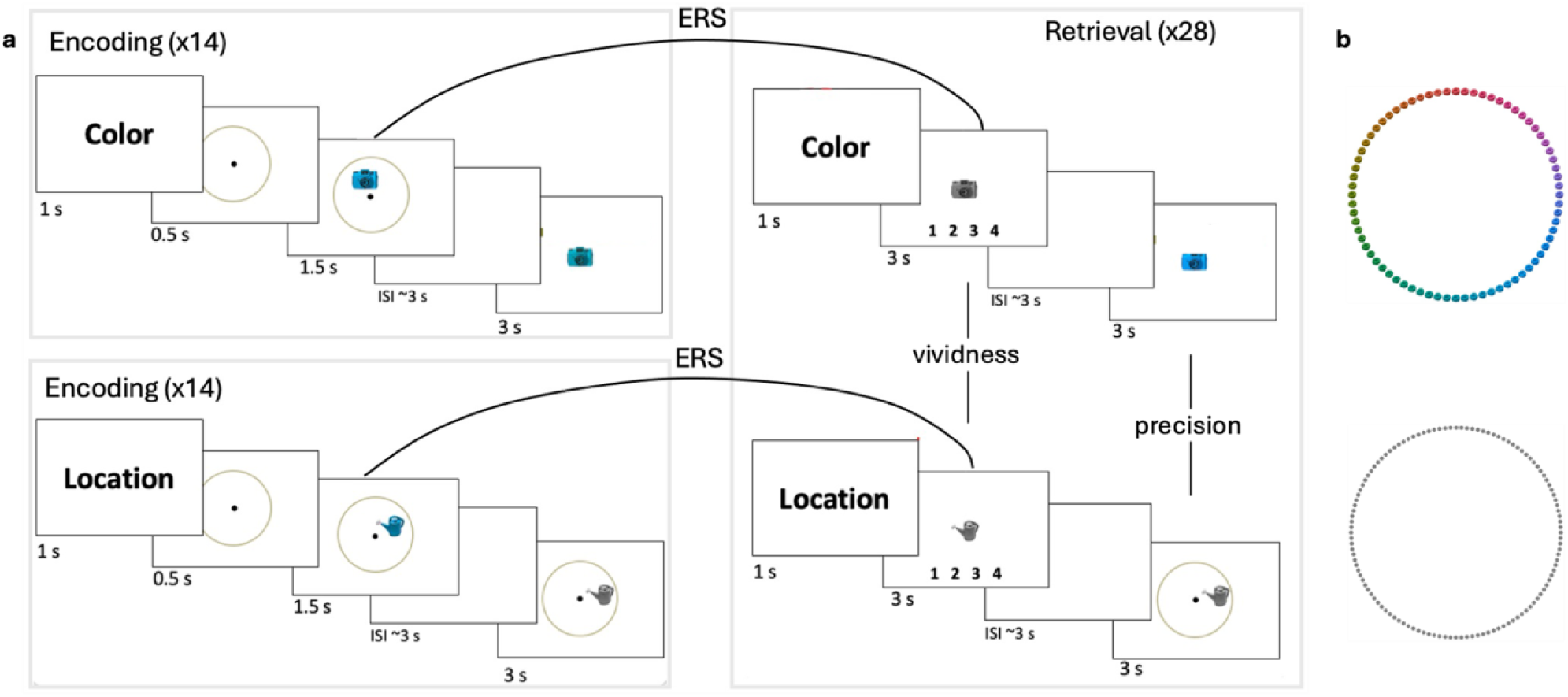
Experimental paradigm. (**a**) During encoding, participants were asked to reproduce the color or location of the object after a varying delay (mean: 3s). Color and location encoding tasks were separated into mini blocks. During retrieval, participants were cued to recall either color or location of the object. Color and location retrieval trials were intermixed, and the orders of trials were randomized. (**b**) The color and location of each stimulus were sampled from a circular feature space. Note: the color wheel was not shown during the encoding response screen to avoid the interference between color and location.

## Materials and Methods

### fMRI experiment

#### Participants

Fifty-one participants (age range: 18-36 years) completed the fMRI experiment. Participants were recruited from Humboldt University Berlin and surrounding communities. All the subjects were right-handed, had normal or corrected-to-normal vision and no history of psychological or psychiatric disorders. Eleven participants were excluded due to chance level memory performance in at least one condition and/or imaging issues (failed fMRIprep preprocessing, susceptibility artefacts, problematic brain mask, mismatched behavioural and imaging runs long interval scanning sessions). As a result, data analysis was performed on the remaining 40 subjects (21 women, Age_Mean±SD_ = 25.5 ± 4.5 years). Written informed consent was obtained from each subject in accordance with the Declaration of Helsinki and a protocol approved by the Ethics Committee of Humboldt University of Berlin.

### Materials

The stimuli were 336 unicolored everyday objects, adapted from Brady et al. (2013). For each participant, each object was randomly assigned one color in a circular color map, and one location in an invisible circle around fixation. To ensure an even distribution, the color and location feature space were first divided into 6 and 8 subspaces, respectively, and 56 colors and 42 locations were randomly chosen from each subspace. Each stimulus object image had a size of approximately 5° of visual angle when placed on the center of the screen.

### Experimental design

Each scanning run contained two encoding blocks and one retrieval block, with 15 seconds rest between blocks. The entire experiment comprised 12 runs, separated into two sessions on two different days, six runs per session. Each day, participants completed a practice session before entering the scanner to familiarize themselves with the continuous report methods for location and color, as well as the general tasks structure of the experiment.

The experimental paradigm is illustrated in **Figure 1**. At the beginning of each encoding block, instructions for each block directed participants to attend and reproduce the location or the color of the objects. During each trial of an encoding block, participants encoded one feature of an object for 1.5 sec (location or color depending on the block), maintained the feature in working memory for a jittered interval (mean: 3 sec; range: 2-6 sec), and then reported the value of the feature along a continuous scale (max 3 sec). The goal of this working memory task was to focus participant’s attention during encoding on either location or color.

After the two counterbalanced encoding blocks (location and color), a retrieval block included location and color trials in random order, indicated by a 1 sec instruction screen at the beginning of each retrieval trial. In each retrieval trial, a target object in greyscale was presented in the center of the screen for 3 sec. During this period, participants were instructed to recall the cued feature of the object (i.e. imagine the object in the original location or color) and rate the vividness of their memory on a scale from 1 (“cannot recall the feature”) to 4 (“highly vivid memory for the feature”). After a jittered interval (mean: 3 sec; range: 2-6 sec), participant had 5 sec to reproduce the feature in their memory as precisely as possible. Participants adjusted the feature displayed on the screen, clockwise or counter-clockwise in the circular space, by pressing two buttons using the index and middle fingers of their right hand. Each button press led to a small (3° for location, 5° for color) change of the value of feature. A third button, pressed by the left hand, was used to confirm their response. Although color values were sampled from a circular parameter space, the color wheel was not shown on the screen to avoid the interference between color and location information. Encoding and retrieval trials were separated by jittered intertrial intervals (mean: 3 sec; range: 2-6 sec). Each stimulus object was assigned to one type of encoding task and one type of retrieval task, and the assignments of objects to encoding-retrieval task combinations was randomized across participants. There were four encoding-retrieval task combinations (location-location, color-color, location-color, color-location), but only trials from the two conditions in which encoding and retrieval tasks matched (location-location and color-color) were analysed in this study. The comparison between matching and mismatching conditions will be the focus of a separate study. The the aim of the current study was to investigate the relationship between neural reinstatement and memory performance of specific features. Therefore, to ensure the same feature was processed during both encoding and retrieval, analysis was restricted to these matched conditions.

### MRI acquisition and brain image preprocessing

All scanning sessions were conducted at the Berlin Center of Advanced Neuroimaging at Charité – Universitätsmedizin Berlin. Structural and functional MR images were collected using a Siemens Magnetom Prisma 3 T scanner (Erlangen, Germany) with a standard 64-channel head coil. A high-resolution structural image was acquired at the beginning of each session using a 3D T1-weighted magnetization prepared gradient-echo (MPRAGE) sequence (208 slices, FoV = 256 mm, voxel size = 0.8 mm^3^, acquisition matrix = 240 x 256 x 167, TR = 2500 ms; TE = 2.22 ms; TI = 1000 ms, flip angle = 8°). Multiband functional images were acquired using a T2*-weighted echo planar imaging (EPI) sequence (72 slices, FoV = 208 mm, voxel size = 2.0 mm^3^, acquisition matrix = 208 × 208 × 144, TR = 800 ms, TE = 37 ms, flip angle = 52°, multi-band acceleration factor = 8). A spin echo field map was acquired to account for the B0 inhomogeneities (72 slices, FoV = 208 mm, acquisition matrix = 208 × 208 × 144, voxel size = 2.0 mm^3^, TR = 8000 ms, TE = 66 ms, flip angle = 90°). Behavioral responses were collected using a 2 x 2 fiber optic response box. MRI-compatible lenses were used to correct for participants’ vision when necessary.

MRI data were preprocessed using fMRIPrep 23.2.0 (Esteban et al., 2019). The T1-weighted volumes were first averaged into a single reference image. The single anatomical reference image then underwent skull-stripping, brain tissue segmentation, surface reconstruction and spatial normalisation. Skull stripping was done by ANT’s brain extraction tool. Brain tissue segmentation was done by FSL’s fast routine. Surface reconstruction was performed with FreeSurfer. Spatial normalization into MNI152 standard space was done by ANT’s antsRegistration scheme. For the preprocessing of the functional images, a non-steady state volume was used as the reference image. This reference image was then contrast-enhanced and skull-stripped. Motion correction of T2*-weighted volumes was done by FSL’s mcflirt routine. Susceptibility distortion correction was done by Phase Encoding POLARity (PEPOLAR) techniques. Co-registration with the T1-weighted image was done by FreeSurfer’s boundary-based registration (bbregister) routine. Finally, the T2*-weighted images were resampled into MNI152 standard space.

### Behavioral data analysis

For each trial during the encoding and episodic retrieval phases, the angular error (0 ± 180°) was calculated as the angular distance between the original feature value and the reported feature value. Considering the continuous nature of the memory report in this experiment, the angular errors were analyzed by fitting a probabilistic mixture model (Bays et al., 2009; Zhang & Luck, 2008) separately for each condition and for each participant. The probabilistic mixture model assumes three types of response errors: a uniform distribution representing random guesses, a von Mises (circular gaussian) distribution centered at the target feature value reflecting memory with various fidelity, and von Mises distributions centered at the values of the non-targets reflecting binding errors. Although both location and color were sampled from circular space, a color wheel was never presented on the screen, and thus, binding errors, which is the confusion between location and color values, were unlikely to occur. Therefore, the probabilistic models were fitted only with the first two components. Hit rate was estimated as the proportion of trials that fell within the von Mises distribution. The standard deviation of the von Mises distribution was also estimated to reflect the memory fidelity. Hit rates of location and color memory were compared using paired-sample t-tests for both encoding and retrieval phase data. Mean vividness of location and color memory were also calculated for each participant and compared using paired-sample t-test. For each participant, Spearman’s correlation between vividness ratings and angular errors was computed separately for location and color memory. The Fisher-transformed Spearman’s correlation across participants was then compared against zero using one-tailed one-sample t-test.

### Encoding-Retrieval Similarity (ERS) Analysis

#### Single trial modeling

To examine neural reinstatement, general linear models (GLMs) were first used to acquire beta estimates for each individual encoding and retrieval trial. A separate GLM was estimated for each trial. Each model included one regressor corresponding to the trial of interest, either encoding or retrieval. Additionally, one regressor modelling all remaining trials of the same phase, one regressor modelling all trials of the opposite phase (encoding or retrieval) and six motion parameters were included as nuisance regressors. In the GLM models, each encoding trial was modeled as a boxcar function with the same onset of the stimulus display and a duration of 1.5 seconds corresponding to the stimulus presentation time. Each retrieval trial was modeled with the onset of the centrally placed greyscale target object and the duration of 3 seconds, corresponding to the period when participants recalled the target feature and rated the memory vividness but before making their continuous report. All regressors were convolved with a canonical hemodynamic response function. Masks for 91 cortical regions of interest (ROIs) and two hippocampal ROIs were extracted from the Harvard-Oxford atlas (Desikan et al., 2006). After GLM estimation, activation patterns were extracted based on the estimated single trial beta, yielding 336 encoding vectors and 336 retrieval vectors for each target ROI. Each vector was z-scored to control for the mean and standard deviation across time to assess the relative activation pattern for each trial within each ROI.

#### Composite memory score

For our first goal of comparing the ERS of distinct perceptual properties of location vs. color, we linked ERS measures to memory using a single composite memory score. This composite score combined the subjective vividness rating and the objective precision measure for each trial. For the precision, angular errors of memory report ranging from -180° to 180° were first converted into precision values using:

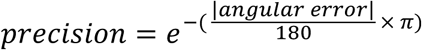

This transformation maps perfect responses (0° error) to a precision value of 1 and maximal errors (±180°) to values near 0. With this non-linear scaling, improvements near perfect recall (e.g., from 10° to 0°) yield a larger increase in precision than improvements near chance (e.g., from 180° to 170°). Vividness ratings and precision values were z-scored within each subject to ensure equal weighting, then summed to form the composite memory score for each trial. Composite memory scores were further divided into four within-subject quartiles to define individualized composite memory score sets (CompMem) for subsequent analysis. Composite memory score sets were defined with individualized standard here to assess the relation between reinstatement and memory at the within-subject level.

#### Item- vs. Set-ERS

ERS scores were calculated as the Pearson correlations between encoding and retrieval vectors, followed by Fisher’s z-transformation. We computed two types of ERS scores: Item-ERS and Set-ERS (**Figure 2**). Item-ERS was calculated between encoding and retrieval trials of the same stimulus item, while Set-ERS was the average of correlations between a retrieval trial and the encoding trials of all other items that share the same feature (location-location or color-color) and same memory score set. The latter allows measuring the effect of Set-ERS on memory performance. Thus, while Set-ERS identifies regions involved in processing location or color in general, Item-ERS identifies regions in which the processing of location or color is sensitive to the particularities of individual stimulus.

**Figure 2.**
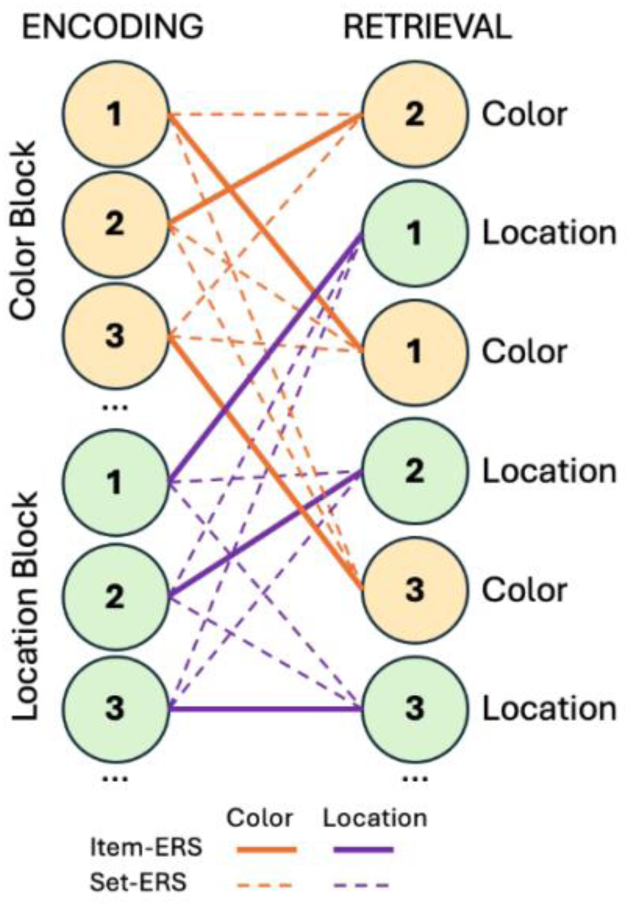
Item- and Set-ERS. ERS analyses for color and location memories were conducted between task-matched encoding and retrieval trials within each run of the experiment. Item-ERS was calculated as neural pattern similarity between encoding and retrieval trials of the same stimulus item. Set-ERS was calculated as neural pattern similarity between encoding and retrieval trials of different stimulus items in the same set. A set includes items that share the same task condition and same memory score (CompMem, vividness or precision category).

Memory score sets were defined differently for the two main analyses. For the location-color comparison, sets were based on quartiles of composite memory scores for location and color within each subject. For the vividness-precision comparison, vividness score sets were given directly by the four-point vividness scale used during retrieval, whereas precision score sets were created by binning continuous precision values into four precision categories (0-0.3, 0.3-0.6, 0.6-0.8, and 0.8-1) to match the four-point scale of vividness. This binning approach yielded comparable distributions across the two measures. Set-ERS (feature-level ERS) pairs were generated within run. Item- and Set-ERS were calculated for each ROI.

#### Linear mixed-effect regression models

To assess differences in reinstatement between location and color memory, we fit a linear mixed-effects model:

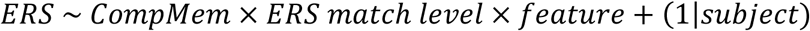

This model predicted ERS scores using fixed terms of CompMem (composite memory score), ERS match level, feature and their interaction terms, as well as random intercepts of subjects. Here, CompMem was a four-level ordered, continuous variable, while ERS match level (Item-ERS/Set-ERS) and feature (location/color) were binary factors. In this model, the main effect of CompMem indicates a memory benefit from overall reinstatement strength regardless of feature. The two-way interaction of CompMem x feature indicates a feature-specific association between reinstatement and memory scores. A three-way interaction, on the other hand, indicates a feature-specific association between trial-specific reinstatement and memory performance.

After identifying a significant main effect of CompMem, we further tested whether this effect truly reflected contributions to both color and location memory, rather than being driven by a single feature. To do so, we fit separate models for color and location:

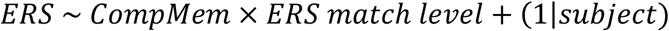

We then identified regions showing a significant main effect of CompMem within each feature and performed a conjunction analysis across the two feature-specific maps. This allowed us to isolate regions in which reinstatement strength contributed to both color and location memory.

To examine differences between subjective vividness and objective precision within each feature memory, separate models were fit for color and locaton trials:

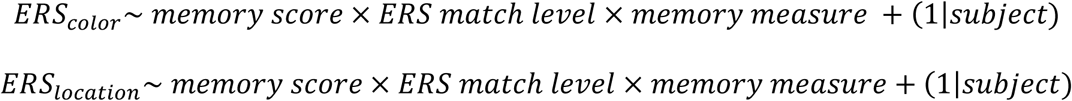

Each model predicts ERS scores using fixed terms of memory score, ERS match level, memory measure and their interaction terms, as well as random intercepts of subjects. Here, memory score was a four-level ordered, continuous variable, while ERS match level (Item-ERS/Set-ERS) and memory measure (precision/vividness) were binary factors. In each model, a significant two-way interaction between memory measure and memory score indicates differences in the relationship between reinstatement and the two memory measures of vividness and precision. A significant three-way interaction indicates the differences in the role of trial-specific reinstatement in supporting vividness vs. precision.

All models were implemented using the “lmer4” package (Bates et al., 2015) in R (version 4.4.3). Given the large number of ROIs, significance of effects was first thresholded by p < 0.05, and then the robustness of which was tested using bootstrapping methods.

#### Bootstrapping procedure

To assess the robustness of observed ERS effects, a non-parametric bootstrapping procedure was done by resampling the data across all participants with replacement for 1000 iterations. In each model, for each ROI showing a significant effect, the effect coefficients were re-estimated in every iteration to generate empirical sampling distributions. The 95% confidence interval (CI) of distribution was then extracted for each effect. In the model comparing location and color, location and color were coded as +0.5 and -0.5, respectively. Therefore, an ROI was considered to show a robust location > color effect if the CI’s lower bound was greater than zero, and a robust color > location effect if the CI’s upper bound was below zero. In the model comparing precision and vividness, precision and vividness were coded as +0.5 and -0.5 in the model, respectively. Therefore, an ROI was considered to show a robust precision > vividness effect if the lower bounds of the 95% CI exceeded zero, and a robust vividness > precision effect if the upper bounds of the 95% CI was smaller than zero.

### Normative study of color naming

To investigate the influence of the linguistic properties of color hues on color memory, 40 native German speakers (age range: 18-36 years, 11 women, Age_Mean±SD_ = 27.8 ± 5.0 years) were recruited and completed a color naming task. Participants were recruited from Prolific (www.prolific.com). In the color naming task, on each trial, participants saw an everyday object (cushion/pitcher/gift bow/origami) and were instructed to type the name of its color. Specifically, participants were allowed to use the same color name for different hues. They were encouraged to provide more specific color name when possible (e.g., silver gray), but they could also use more general color names (e.g., gray) if they felt those were most appropriate. Each participant completed 72 trials, corresponding to the 72 color hues used in the fMRI experiment. The order of the color hues was randomised for each participant. The task was self-paced. Participants submitted their response by pressing a button to move to the next trial.

Responses were minimally cleaned to correct typing errors, and all participants data were pooled. The color names provided for each hue were visualized in **Figure 6a** in the scatter plot. The frequency of each color name provided across the color space were revealed in the line plot. To aid visualization, we applied a smoothing kernel using a sliding window (window size = 3), which reduces the noise from small fluctuations. To test the effect of linguistic categories on color memory, the bias of episodic memory for each hue was calculated based on the pooled data from the main experiment. The direction of the bias was determined by the predominant direction of errors, and the size of bias was calculated as the ratio between errors in the predominant vs. non-predominant direction.

To assess the role of linguistic processing in color memory quality, we calculated color name richness, defined as the number of different names provided for each hue. We fit linear mixed-effects models:

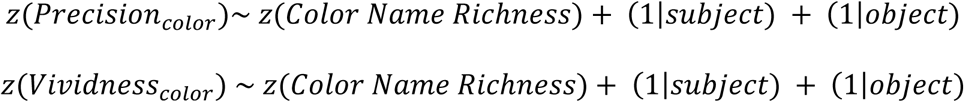

where precision and vividness were continuous variables measured during the fMRI experiment. The main effect of color name richness reflects whether color memory, either objective precision or subjective vividness, benefits from more potential labels. In the first two models, only precision showed a reliable relationship with color name richness, whereas vividness did not. Given this pattern, the following moderation analysis examining ERS was conducted using precision, as it was the only memory outcome demonstrating sensitivity to linguistic factors.

Finally, to test whether linguistic processing moderates how reinstatement supports color memory, we fit a linear mixed-effects model:

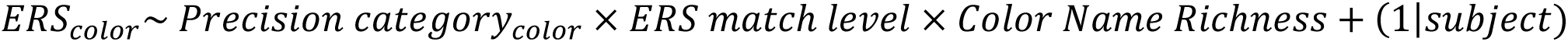

This model predicts ERS scores during color memory trials using fixed terms of color memory precision category, ERS match level, color name richness and their interaction terms, as well as random intercepts of subjects. In this model, a significant two-way interaction between color name richness and color memory score would indicate that the relationship between reinstatement and color memory varies with color name richness of the hues. A significant three-way interaction would indicate that trial-specific reinstatement differentially supports color memory depending on its color name richness.

## Results

### Behavioral results of fMRI paradigm

During the encoding phase, we used a working memory task with a short delay (average: 3 sec) not to study working memory process but to ensure that attention would focus on either the location or color of each object. Given the short delay, accuracy was almost perfect (**Figure 3a**). Estimated by the mixed model, accuracy was 0.99 (SD = 0.01) for location, and 0.96 (SD = 0.04) for color. The angular error between reported and true values of the feature had a standard deviation of 10.31 degrees (SD = 2.42) for location and 25.34 degrees (SD = 3.85) for color, significantly larger for color memory (*t*_(39)_ = 24.73, *p* < .001). The high hit rates confirm that the subjects were able to attend to the target feature of the stimuli during encoding.

**Figure 3.**
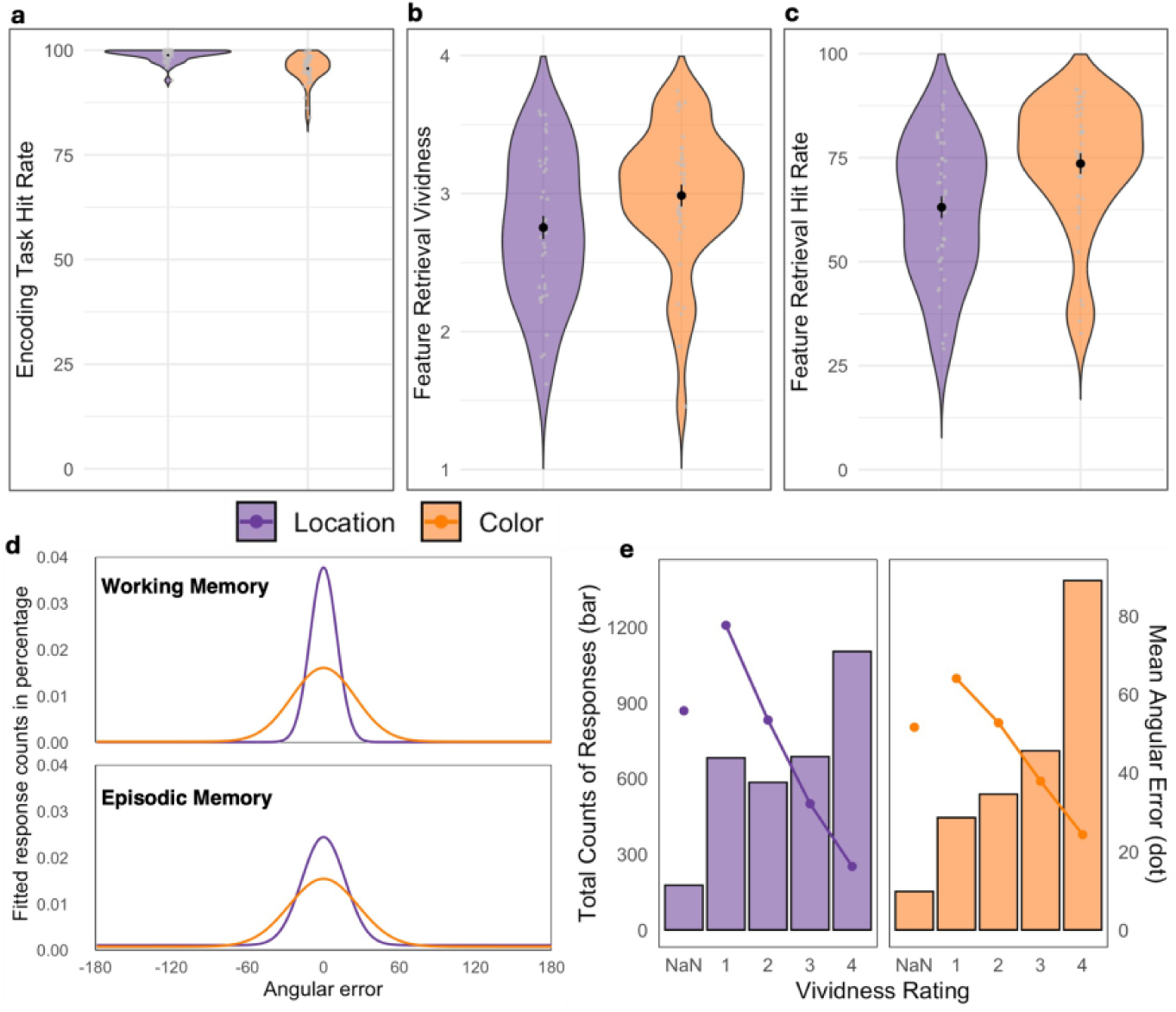
Behavioral results of encoding and retrieval tasks. (**a**) Model-estimated hit rate of color and location working memory during encoding. Black dots and error bars indicate group mean and standard error of the mean (SEM). (**b**) Mean color and location memory vividness during retrieval. Error bars indicate the standard error of the means (SEM). (**c**) Model-estimated hit rate of color and location episodic memory during retrieval. (**d**) Best-fitting model probability density function of aggregated color (orange) and location (purple) errors during encoding task (upper panel) and episodic retrieval (lower panel). Note that episodic memory has an obvious uniform distribution fitted by the model, indicating the guesses among responses. (**e**) Bar plots indicate the total counts of each vividness rating in color and location memory retrieval; line plots indicate the mean angular error corresponding to each vividness rating.

During the retrieval phase, subjects were cued to recall one of the two features of each object. First, they rated the vividness of recalled feature, and then reported the feature value as precisely as possible on a continuous scale. Vividness ratings were higher (*t*_(39)_ = 4.54, *p* < .001) for color (M = 2.99; SD = 0.50) than location memory (M = 2.75; SD = 0.53) (**Figure 3b**). Mean hit rate (defined as the proportion of trials falling within the von Mises distribution; see Behavioral Methods) was also higher (*t*_(39)_ = 3.95, *p* < .001) for color (M = 0.74; SD = 0.16) than location memory (M = 0.63; SD = 0.16) (**Figure 3c**). For hit trials, the angular error (original vs. reported value) had a mean standard deviation of 28.05 degrees (SD = 8.37) for color and 18.78 degrees (SD = 5.84) for location (**Figure 3d**), significantly larger for the color memory (*t*_(39)_ = 6.82, *p* < .001), indicating a higher precision of location memory. Thus, although the proportion of hits (signal vs. noise) was higher for color than location, the precision of these hits (signal exactness) was greater for location than color. This precision difference between location and color was also evident during the encoding task and likely reflects the inherently different resolution with which people can distinguish and report these spatial and chromatic features.

Finally, we investigated the relationship between vividness ratings and precision. As illustrated by **Figure 3e**, there was a moderate negative within-subject Spearman correlation between vividness rating and angular error both for color (ρ = -0.29) and location (ρ = -0.45). Fisher-z transformed correlations were significantly smaller than zero both for color (*t*_(39)_ = -14.33, *p* < .001) and location (*t*_(39)_ = -24.99, *p* < .001). The significant association between vividness and precision indicates that vividness ratings, although subjective, are a valid memory measure.

In sum, memory accuracy was at ceiling during encoding and high during retrieval. During retrieval, both vividness and accuracy were higher for color than for location, although precision was higher for location than color. Finally, for both color and location, vividness ratings and angular errors were negatively related, indicating that vividness is a valid memory measure.

### fMRI results

#### Goal 1: Neural reinstatement for color and location memory

Our first goal was to use ERS to identify the perceptual specificity of neural reinstatement for location and color memory. To address this goal, we used a linear mixed-effects model, focusing explaining the variance in ROI-level ERS as predicted by (1) the main effect of the composite memory score (CompMem), (2) the interaction between CompMem and stimulus Feature (location vs. color), and (3) the three-way interaction between CompMem, ERS level (Item- vs. Set-ERS), and Feature. This model was applied independently at each region of interest (ROI), and the robustness of the observed effects was verified using a bootstrapping procedure. The results are shown in **Table 1** and **Figure 4**.

**Figure 4.**
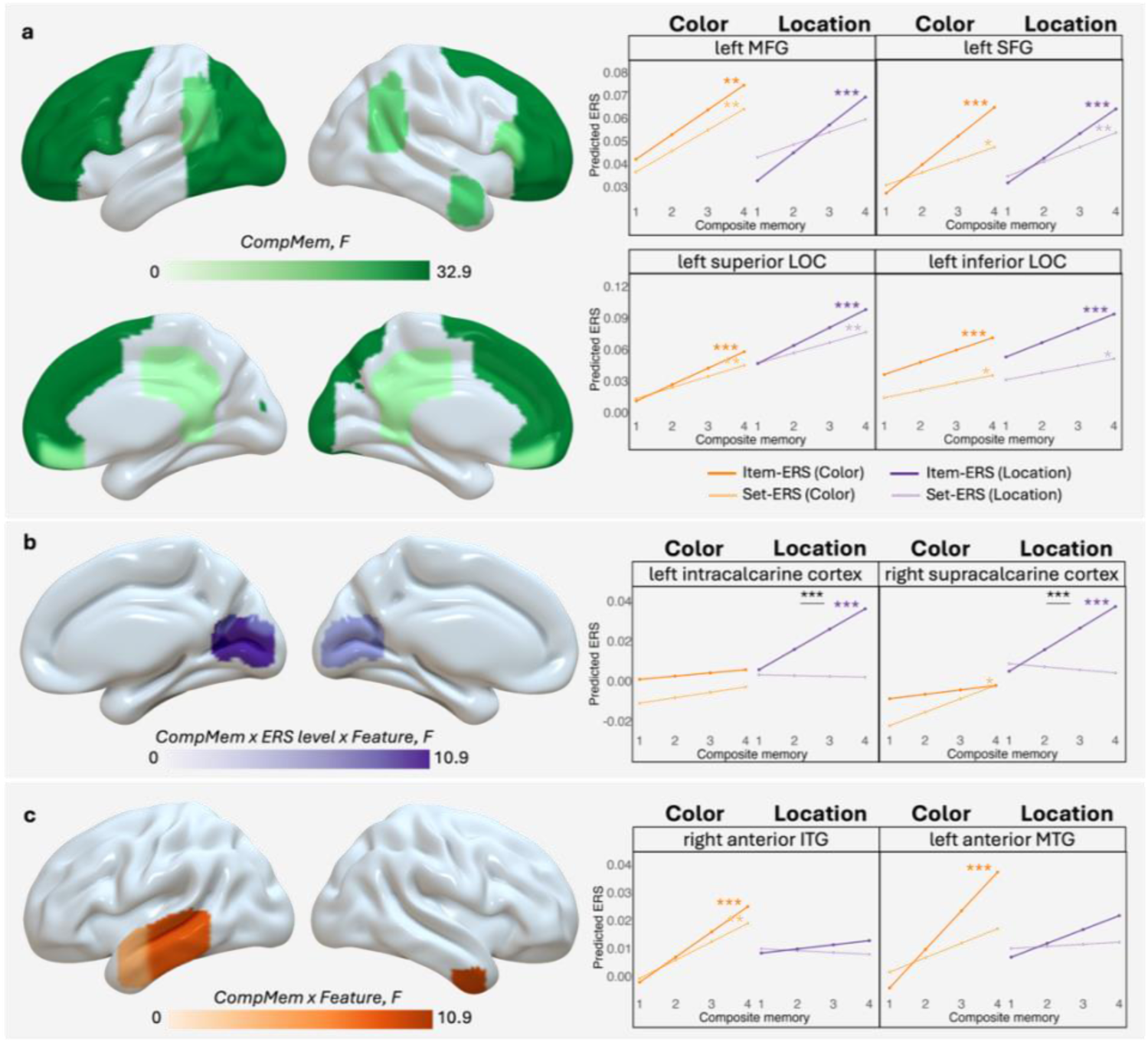
Memory-enhancing reinstatement regions differentiating for color vs. location. (**a**) Regions where feature-level reinstatement was associated with memory performance regardless of color or location. (**b**) Regions where trial-specific reinstatement was associated with location (purple) > color memory. No regions were found with opposite effect. (**c**) Regions where overall reinstatement was associated with color (orange) > location memory. No regions were found with opposite effect. [Line plots are illustrations of representative regions for each effect. Colored asterisks next to each line indicates a significant correlation between ERS and memory score at the corresponding ERS-level for the corresponding feature. Black asterisks at the top of each panel indicate a significant interaction between ERS-level and memory score for the corresponding feature. *p<.05, **p<.01, ***p<.001 (uncorrected). All the regions plotted achieved significance in the bootstrapping test. MFG: middle frontal gyrus; SFG: superior frontal gyrus; LOC: lateral occipital cortex; ITG: inferior temporal gyrus; MTG: middle temporal gyrus.]

**Table 1.** Regions showing reinstatement-memory related effects.

| Region | Hemi | Denominator<br>DF | F | uncorrected<br>p |
| --- | --- | --- | --- | --- |
| **Main effect of CompMem** |  |  |  |  |
| sLOC | L | 9688.041 | 60.818 | .000 |
| IFG_tri | L | 9684.845 | 46.914 | .000 |
| FP | L | 9701.917 | 45.910 | .000 |
| SFG | L | 9677.803 | 44.369 | .000 |
| SFG | R | 9701.994 | 42.095 | .000 |
| iLOC | L | 9677.710 | 39.825 | .000 |
| toMTG | L | 9700.801 | 37.419 | .000 |
| FP | R | 9701.437 | 34.909 | .000 |
| PaCiG | R | 9698.838 | 33.602 | .000 |
| MidFG | L | 9676.590 | 33.373 | .000 |
| toITG | L | 9694.444 | 32.246 | .000 |
| OP | L | 9678.634 | 32.075 | .000 |
| IFG_oper | L | 9679.517 | 31.642 | .000 |
| pMTG † | L | 9701.233 | 31.485 | .000 |
| PaCiG | L | 9685.549 | 31.274 | .000 |
| AG | L | 9699.221 | 25.368 | .000 |
| aMTG | R | 9699.753 | 24.495 | .000 |
| plTG | L | 9690.281 | 24.479 | .000 |
| AG | R | 9696.239 | 22.093 | .000 |
| pSMG | L | 9688.568 | 21.878 | .000 |
| IFG_tri | R | 9699.827 | 20.721 | .000 |
| MedFC |  | 9681.585 | 19.081 | .000 |
| iLOC | R | 9676.569 | 18.638 | .000 |
| aMTG † | L | 9697.289 | 15.538 | .000 |
| PC |  | 9700.160 | 15.098 | .000 |
| pMTG | R | 9701.862 | 14.204 | .000 |
| alTG | L | 9427.762 | 13.203 | .000 |
| FOrb | L | 9700.097 | 12.712 | .000 |
| alTG † | R | 9701.858 | 12.680 | .000 |
| **Main effect of CompMem (continue)** |  |  |  |  |
| sLOC | R | 9697.009 | 12.526 | .000 |
| ICC ‡ | R | 9689.947 | 11.029 | .001 |
| pSTG | L | 9696.746 | 10.339 | .001 |
| pTFusC | L | 9691.281 | 10.202 | .001 |
| ICC ‡ | L | 9685.755 | 10.073 | .002 |
| toMTG | R | 9700.503 | 9.972 | .002 |
| aSMG | L | 9683.698 | 9.754 | .002 |
| OFusG | L | 9680.776 | 9.563 | .002 |
| SCC ‡ | R | 9693.957 | 9.531 | .002 |
| MidFG | R | 9701.981 | 9.442 | .002 |
| OP | R | 9677.928 | 9.291 | .002 |
| SCC ‡ | L | 9690.369 | 9.288 | .002 |
| pSMG | R | 9687.928 | 9.198 | .002 |
| pSTG | R | 9699.918 | 8.307 | .004 |
| TOFusC | L | 9686.717 | 8.226 | .004 |
| TP | L | 9701.246 | 8.123 | .004 |
| FO | L | 9692.718 | 7.420 | .006 |
| FOrb | R | 9649.159 | 6.260 | .012 |
| aSTG | R | 9683.363 | 5.707 | .017 |
| **CompMem x feature interaction: color > location** |  |  |  |  |
| alTG | R | 9045.633 | 10.012 | .002 |
| pMTG | L | 9300.051 | 8.750 | .003 |
| aMTG | L | 8593.878 | 4.402 | .036 |
| **CompMem x ERS_level x feature interaction: location > color** |  |  |  |  |
| ICC | R | 9662.670 | 14.760 | .000 |
| SCC | R | 9663.359 | 8.243 | .004 |
| ICC | L | 9663.055 | 6.949 | .008 |
| SCC | L | 9663.027 | 6.140 | .013 |

##### Main effect of CompMem

The main effect of CompMem revealed regions in which overall reinstatement was predicted by memory performance (see Table 1). These areas do not distinguish between memory-related reinstatement that are specific to a perceptual property (color or location) or are sensitive to trial-specific information. To ensure that the effects are truly common for color and location, we identified effects of CompMem separately for color and location and performed a conjunction analysis between them. The regions that survived the conjunction analysis are highlighted in Table 1 and shown in **Figure 4a**. These are likely regions that contribute to visual memory in general. Consistent with this idea, they included frontoparietal areas associated with attentional control processes (Eichenbaum, 2017; Sestieri et al., 2017) and posterior visual areas linked to visual processing (Bone et al., 2020).

##### CompMem x Feature interaction

The interaction revealed regions where reinstatement contributed differently to CompMem for color and location. **Table 1** and **Figure 4b** show regions where ERS was associated more strongly with color relative to location memory. As illustrated by **Figure 4b**, these regions include the left middle temporal gyrus and right anterior inferior temporal gyrus (aMTG_l: *F*_(1, 8593.88)_ = 4.40, *p* = .036; pMTG_l: *F*_(1, 9300.05)_ = 8.75, *p* = .003; aITG_r: *F*_(1, 9045.63)_ = 7.12, *p* = .002). No regions demonstrating an interaction showed the opposite pattern, i.e., location > color. As illustrated by the graphs in **Figure 4c**, for color, ERS increased as a function of CompMem, whereas for location, ERS was largely flat. Given the role of lateral and anterior temporal regions in semantic and linguistic processes, it is possible that these functions contribute to memory for color more than for location.

##### CompMem x ERS match level x Feature interaction

The three-way interaction identified regions where trial-specific reinstatement differentially relates to color and location memory performance. This three-way interaction was significant in early visual cortex, particularly in bilateral calcarine cortex, where the association between trial-specific reinstatement and CompMem was greater for location than for color (ICC_r: *F*_(1, 9662.67)_ = 14.76, *p* < .001; ICC_l: *F*_(1, 9663.06)_ = 6.95, *p* = .008) (see **Figure 4b**). No regions showed the reversed pattern (color > location). As shown by the graphs in Figure 4C, in early visual cortex, CompMem scores for location increased with Item-ERS but not for Set-ERS. These results suggest early visual cortex stores trial-specific location representations, but not color information, and that this detailed location information is reactivated during retrieval, driving memory performance.

#### Goal 2: Neural reinstatement for vividness and precision

Our second goal was to compare neural reinstatement supporting subjective vividness and objective precision. For this goal, we ran separate models for color and location, examining (1) the two-way interaction between memory scores (1 to 4) and memory measure (vividness vs. precision), and (2) the three-way interaction between memory scores, ERS match level and memory measure. Results are shown in **Figure 5** and **Table 2**.

**Figure 5.**
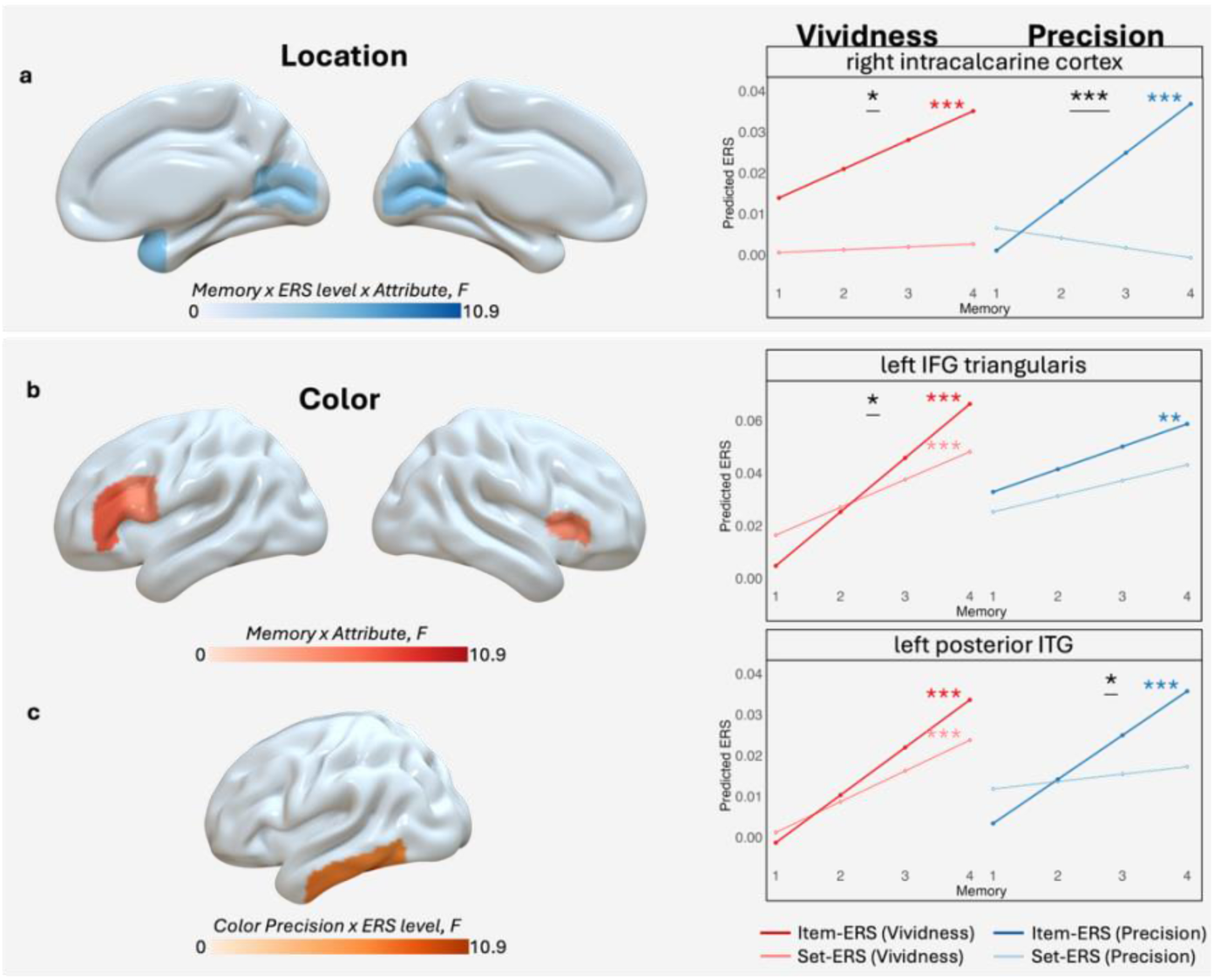
Effects of memory scores and memory measure (vividness vs. precision), separately for color and location memory. (**a**) Regions where trial-specific reinstatement was associated with location precision (blue) more than vividness (red), among which right intracalcarine cortex is illustrated as a representative region. No regions showing three-way interaction were found with the opposite effect. (**b**) Regions where overall reinstatement was associated with color vividness (red) more than precision (blue), among which left IFG triangularis is illustrated as a representative region. No regions showing two-way interaction were found with the opposite effect. (**c**) Regions showing trial-specific reinstatement contribution to color precision. Significant regions are left inferior temporal gyrus (ITG). [Colored asterisks next to each line indicates a significant correlation between ERS and memory score at the corresponding ERS-level for the corresponding measure. Black asterisks at the top of each panel indicate a significant interaction between ERS-level and memory score for the corresponding measure. *p<.05, **p<.01, ***p<.001 (uncorrected). All the regions plotted achieved significance in the bootstrapping test. IFG: inferior frontal gyrus.]

**Figure 6.**
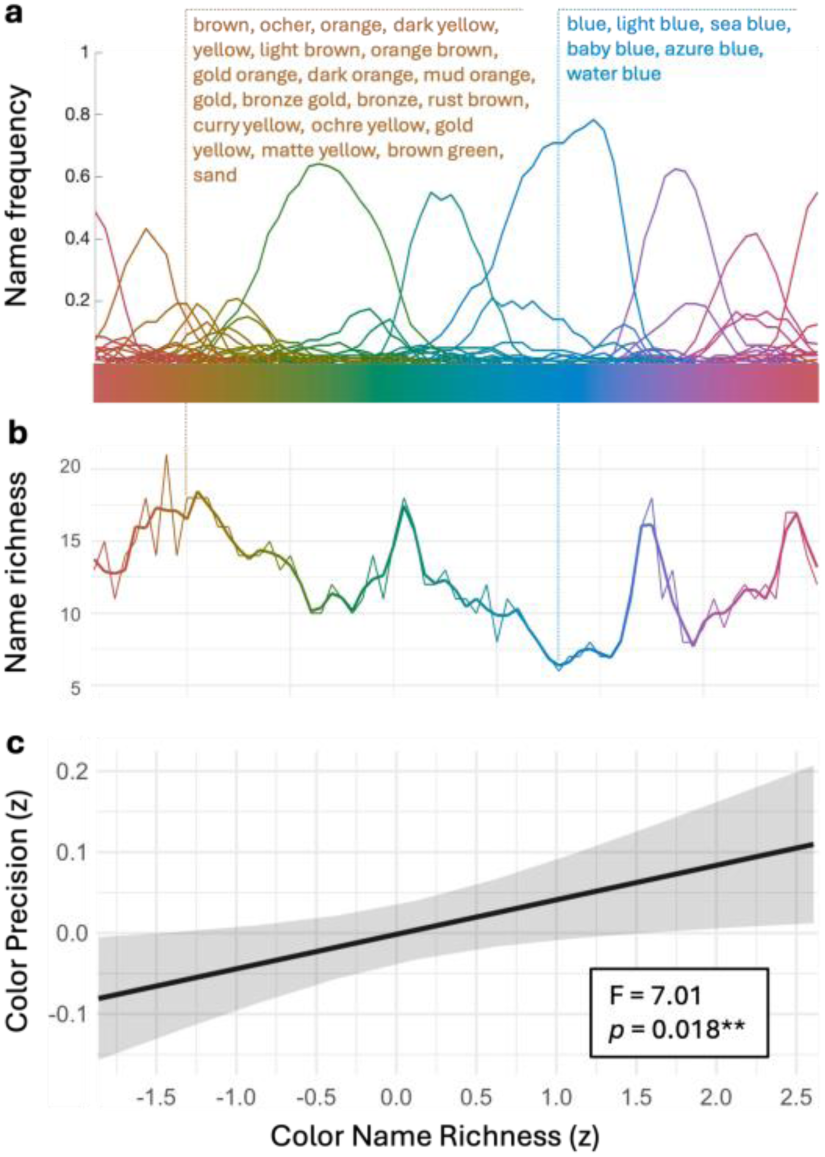
Linguistic effects on color memory. (**a**) Distribution of color names across color hues. Names for two example hues are listed to demonstrate the difference in color name richness. (**b**) Distribution of color name richness across color hues. Here, color name richness is defined by different color names provided for a given color hue by a separate group of 40 subjects. (**c**) Color memory significantly increased with color name richness.

**Table 2.** Regions showing vividness vs. precision contrast for each feature memory. [Abbreviations: IFG_tri: inferior frontal gyrus triangularis; FO: frontal operculum; IFG_oper: inferior frontal gyrus opercularis; TP: temporal pole; ICC: intracalcarine cortex; PP: planum polare; SCC: supracalcarine cortex.]

| Region | Hemi | Denominator<br>DF | F | uncorrected<br>p |
| --- | --- | --- | --- | --- |
| **Location memory x ERS_level x Attribute interaction: precision > vividness** |  |  |  |  |
| TP | R | 9766.088 | 5.175 | .023 |
| ICC | L | 9767.040 | 5.037 | .025 |
| PP | R | 9765.965 | 4.368 | .037 |
| ICC | R | 9766.896 | 4.153 | .042 |
| SCC | L | 9766.503 | 3.979 | .046 |
| **Color memory x Attribute interaction: vividness > precision** |  |  |  |  |
| IFG_tri | L | 10031.716 | 6.844 | .009 |
| FO | R | 9521.374 | 4.830 | .028 |
| IFG_oper | L | 10035.385 | 4.800 | .028 |
| FO | L | 9892.571 | 4.024 | .045 |

##### Vividness vs. precision in location memory

For location memory, a significant three-way interaction revealed brain regions where trial-specific reinstatement differentially contributed to vividness and precision. This effect was only observed for location precision > vividness. **Figure 5a** shows that in the bilateral calcarine cortex, trial-specific reinstatement was more strongly associated with location precision than with vividness (ICC_l: *F*_(1, 9767.04)_ = 5.04, *p* = .025; ICC_r: *F*_(1, 9766.90)_ = 4.15, *p* = .042). Given the role of calcarine cortex in early visual processing, these results suggest that the reinstatement of fine-grained sensory detail benefits the objective precision of location memory more than its subjective vividness. Note, however, that trial-specific reinstatement in early visual cortex also predicted location vividness, albeit to a lesser extent. No ROI showed a significant two-way interaction between memory score and memory measure.

##### Vividness vs. precision in color memory

For color memory, a significant interaction between memory measure and memory score revealed regions where reinstatement differentially contributed to its vividness vs. precision. This effect was observed only for color vividness > precision. **Figure 5b** shows that for bilateral inferior frontal gyrus, reinstatement is associate more strongly with color vividness than precision (IFG_tri_l: *F*_(1, 10031.72)_ = 6.84, *p* = .009; IFG_oper_l: *F*_(1, 10035.39)_ = 4.80, *p* = .028). Considering that the IFG has been consistently implicated in controlled semantic activation (Jackson, 2021; Martin & Cheng, 2006), these findings suggest that semantic activation of color, e.g., “blueish”, may give a stronger feeling of color vividness more than contributing to its objective precision.

*Post-hoc analysis.* Although no ROI showed a significant three-way interaction (memory score × ERS match level × measure), we reasoned that trial-specific reinstatement should nevertheless contribute to color precision. A *post hoc* analysis confirmed that trial-specific reinstatement in the posterior inferior temporal gyrus was associated with higher color precision (pITG_l: *F*_(1, 4908.23)_ = 7.58, *p* = 0.006; toITG_l: *F*_(1, 4907.78)_ = 4.96, *p* = 0.026) (**Figure 5c**). This area, encompassing the classic color-selective area V4, is known to represent color in a geometry consistent with human perceptual similarity (Brouwer & Heeger, 2009). Thus, while frontal reinstatement seems to enhance the vividness of color memory via semantic activation, precise color memory may rely more on trial-specific reinstatement in perceptual regions.

#### Goal 3: Linguistic effect on color reinstatement

Our third goal was to examine whether linguistic labelling shapes color memory reinstatement. First, we plotted the results of the color naming experiment, in which participants named objects in 72 different color hues. As illustrated by **Figure 6a**, basic color categories like “red”, “orange”, “green”, “turquoise”, “blue”, “purple” and “pink” emerged. However, our goal was not to identify color categories. **Figure 6a** shows a particular yellowish hue described with 18 different names and a bluish hue described with only 6 names. This illustrates that the richness of language used to describe each color varies across the spectrum.

To capture this systematically, we measured *color name richness* by calculating the number of different words participants used to describe each color. **Figure 6b** displays color name richness across the color spectrum. The color spectrum regions with greater color name richness (e.g. over 15 different names for each hue) are the areas between red and green, between green and turquoise, between blue and purple, and between pink and red. These regions correspond to areas with multiple overlapping distributions in **Figure 6a**.

Next, we measured the effect of color name richness on memory-based color precision in the main study. As shown in **Figure 6c**, color name richness was associated with greater color precision (*F*_(1, 3082.62)_ = 5.64, *p* = 0.018). This finding is consistent with the previous work from (Hasantash & Afraz, 2020). One possible interpretation is that name richness reflects the availability of multiple categorical anchors, and such linguistic scaffolding helps bind continuous perceptual inputs to fine-grained lexical representations, thereby supporting color memory. Finally, to directly test our interpretation that the stronger reinstatement effect for color memory in the temporal gyrus reflects linguistic processing, we examined whether reinstatement in these regions is moderated by linguistic properties of the colors. We fit a linear mixed-effects model, predicting ERS scores using color name richness, reinstatement level, and memory precision. This analysis revealed a significant three-way interaction in the right anterior inferior temporal gyrus (aITG) (*F*_(1, 4603.40)_ = 4.91, *p* = 0.027). To illustrate this effect, **Figure 7** plots the relationship between color precision and trial-specific ERS (the difference between Item- and Stimulus-independent ERS), with hues split into high versus low color name richness by a median split. As shown, color precision was positively associated with trial-specific ERS, and this association was specific for hues with high color name richness. These results suggested that the right aITG supports color precision through trial-specific reinstatement, but such reinstatement is facilitated when colors can be richly and flexibly labelled with language. Together, the color naming task demonstrates that linguistic properties play a functional role in color memory and moderate its neural reinstatement.

**Figure 7.**
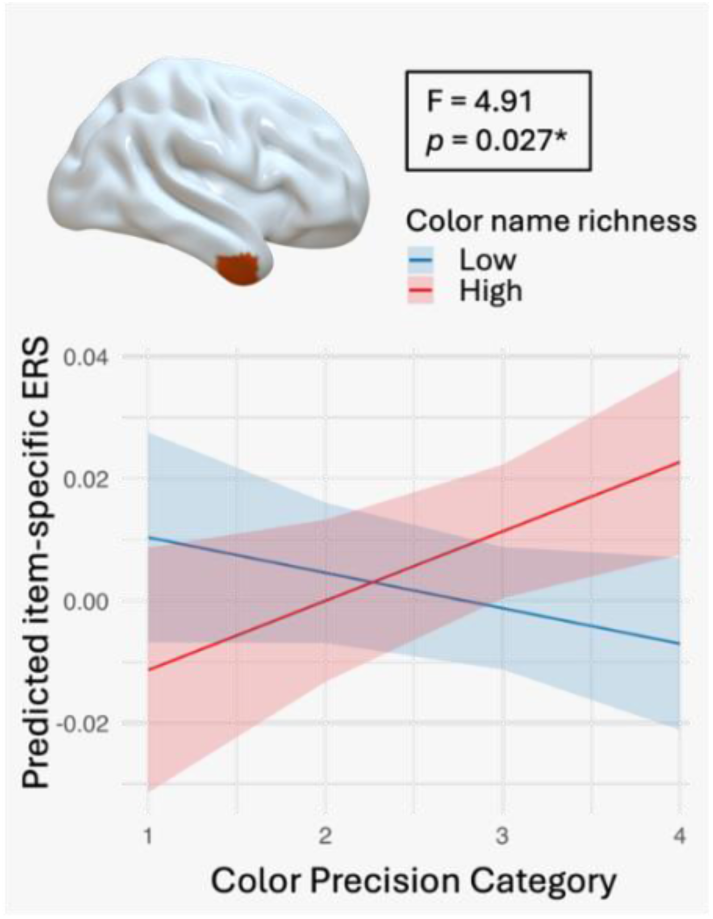
Linguistic moderation of color reinstatement. Relation between right inferior temporal gyrus (aITG_r) reinstatement and color memory is moderated by color name richness. Particularly, trial-specific reinstatement contributes to color memory more for the colors with high name richness.

## Discussion

Although reinstatement is central to episodic memory, available evidence is unclear on how it differs for various perceptual features, supports subjective versus objective memory, and relies on linguistic information. Using encoding–retrieval similarity (ERS) measures, this study yielded three main sets of findings. First, reinstatement in a distributed frontoparietal-visual network supported memory for both features, with early visual cortex preferentially supporting location and temporal cortices preferentially supporting color (**Figure 4**). Second, early visual cortex reinstatement contributed to location precision and vividness, whereas reinstatement in the inferior frontal gyrus and posterior inferior temporal regions contributed to color vividness and color precision, respectively (**Figure 5**). Finally, the number of labels a particular color can have (color name richness) enhanced color precision and modulated trial-specific color reinstatement in right anterior inferior temporal gyrus (**Figure 7**).

### Location vs. color memory reinstatement

Our first goal was to compare neural reinstatement for location vs. color memory, and both similarities and differences were found. A key similarity was feature-level reinstatement in frontoparietal regions and lateral occipital cortex (LOC), which supported memory for both features (**Figure 4a**). This shared reinstatement likely reflects domain-general processes that enhance memory encoding and retrieval regardless of feature type. The answer is likely different across regions. In frontoparietal areas, it most likely reflects attentional and/or control processes (Ray et al., 2020) that are known to facilitate episodic memory (Greene & Naveh-Benjamin, 2022; Madore & Wagner, 2022). Consistent with this interpretation, neuroimaging studies have shown that frontoparietal activity is associated with both successful encoding (Xue et al., 2013) and retrieval (Iidaka et al., 2006; Kwok & Macaluso, 2015), with overlap between encoding- and retrieval-related regions (Kim et al., 2010). In contrast, reinstatement in LOC likely reflects visual object processing, which is critical for object-based memory (Guo & Yang, 2023; Karanian & Slotnick, 2015). Together, these findings suggest that attentional control and visual processes are required for successful encoding and retrieval, accounting for the broad reinstatement effects in frontoparietal and LOC regions.

In contrast to these shared regions, other brain areas were differentially involved in location vs. color memory. Based on evidence for dual visual pathways (Ungerleider & Mishkin, 1982), we predicted dorsal pathway reinstatement for location and ventral pathway reinstatement for color memory. Yet, we found feature-specific reinstatement in other regions. Location memory was selectively associated with trial-specific reinstatement in early visual cortex (**Figure 4b**). Although this region codes both spatial and color information (Vo et al., 2022; Yan et al., 2023), its preferential link to location memory might reflect its retinotopic organization (Hubel & Wiesel, 1959; Sereno et al., 1995), which was observed not only during visual perception but also during visual imagery and recall (Slotnick, 2009; Slotnick et al., 2005; Vicente-Grabovetsky et al., 2014). Even though we did not control eye movements, seeing an object at one location during encoding and imagining it at the same location during retrieval likely recruited similar retinotopic areas, producing the observed memory-enhancing effect.

Finally, several regions showed greater reinstatement for color than for location. Color memory was preferentially associated with feature-level reinstatement in the left MTG and right anterior ITG. (**Figure 4c**). This finding aligns with evidence that lesions or stimulation in these regions affect color discrimination (Buckley et al., 1997), color naming (Roux et al., 2006), and color memory capacity (Chiou & Lambon Ralph, 2018). Posterior MTG has been implicated in receptive semantic control (Davey et al., 2016; Noonan et al., 2013), and although both anterior temporal lobes function as semantic hubs (Ralph et al., 2017), the right anterior temporal lobe was more specific for perceptual inputs (Rice et al., 2015). Compared to location, color is more tightly linked to semantic knowledge (Guilbeault et al., 2020) and benefits more from semantic processing (Heurley et al., 2013a). Therefore, semantic processing might underlie the preferential involvement of MTG and ITG in color memory.

### Subjective vividness vs. objective precision

Our second goal was to compare neural reinstatement supporting subjective vividness vs. objective precision. For location memory, trial-specific reinstatement in early visual cortex was tied to its precision more than its vividness (**Figure 5a**). This finding is consistent with evidence linking early visual reinstatement to precise orientation memory (Ester et al., 2013). Although no region showed a stronger association with location vividness than precision, location vividness was nevertheless also supported by trial-specific reinstatement in early visual cortex (**Figure 5a**). The absence of vividness-specific reinstatement for location may attribute to that spatial location per se does not vary in visual and semantic properties, factors contributing to memory vividness of complex images (Morales-Torres et al., 2025). Together with the moderate behavioral correlation between location precision and vividness (**Figure 3e**), these findings suggests that vividness judgments for location might be informed by the same sensory reinstatement mechanisms that support its precise recall. For color, memory precision was supported by trial-specific reinstatement in the posterior inferior temporal gyrus (pITG) (**Figure 5c**), which includes area V4, a region strongly associated with color processing (Zeki, 1973). In humans, V4 lesions can cause achromatopsia and impair color constancy, linking this region to the conscious experience of color (Cowey & Heywood, 1997; Zeki, 1998). Multivariate fMRI studies have shown that human V4 codes for both perceived and imagined colors (Bannert & Bartels, 2018; Brouwer & Heeger, 2009), which explains why it can be engaged during both encoding and retrieval of colored objects and displayed significant reinstatement. In addition to pITG, trial-specific reinstatement in the right anterior ITG also contributed to color precision. As noted, this region is associated with perceptual semantics (Rice et al., 2015), indicating that both perceptual and semantic representations support precise color memory.

In contrast, feature-level reinstatement in bilateral inferior frontal gyrus (IFG) was more strongly associated with color vividness than precision (**Figure 5b**). The left IFG, including Broca’s area, is implicated in semantic retrieval (Foundas et al., 1998; Liakakis et al., 2011), particularly under high retrieval demands (Martin & Cheng, 2006; Whitney et al., 2011). The right IFG has been similarly linked to covert naming (Rivas-Fernández et al., 2021) and lexical selection (Vigneau et al., 2011). These findings suggest that vivid color memory may be enriched by semantic activation in IFG, consistent with evidence that semantic information can enrich subjective vividness of visual recollection (Cooper & Ritchey, 2022; Morales-Torres et al., 2025).

### Role of language in processing and reinstatement

Our third goal was to examine how language shapes color memory reinstatement. In a behavioral color naming experiment, we measured the number of labels available for each hue across the color spectrum. As expected, we found that greater color name richness predicted higher color memory precision (**Figure 6c**), consistent with previous evidence that verbal labels enhance color memory (Hasantash & Afraz, 2020). Critically, color name richness moderated reinstatement in the right aITG (**Figure 7**): trial-specific reinstatement in this region supported color precision only for hues with higher name richness. Given evidence that linguistic processing modulates color perception (Heurley et al., 2013b), the right aITG might enhance color memory precision through similar linguistic scaffolding of color perception and recall.

This finding suggests that the way reinstatement mediates memory for simple features depends on the degree to which the feature is supported by verbal scaffolds. For less verbalizable feature, like spatial location, memory relies mainly on low-level sensory reinstatement, consistent with findings for orientation memory, another basic visual feature with limited linguistic coding (Ester et al., 2013). In contrast, for features that are readily verbalizable, such as color, memory benefits from both perceptual and linguistic/semantic reinstatement. It should be acknowledged, however, that visual-spatial memory may also be affected by language (Pereira Seabra et al., 2025), suggesting that linguistic coding may broadly shape visual memory, not just for color.

More broadly, our results highlight a dynamic relationship between reinstatement, precision and vividness. When verbalization is limited, sensory reinstatement primarily supports memory precision and likely informs vividness judgement. When verbal coding is easier, linguistically-mediated perceptual reinstatement continues to support objective precision, while subjective vividness can be more influenced by the general IFG reinstatement, likely reflecting semantic activation, in line with evidence that memory vividness depends on semantic gist recalled (Baddeley & Andrade, 2000; Cooper & Ritchey, 2022) or semantic distinctiveness of stimuli (Morales-Torres et al., 2025). Overall, perceptual and linguistic reinstatement may complement each other in supporting visual memory, with their relative contributions determined by the representational format and lexical richness of the memory content.

## Conclusion

In summary, our findings show that neural reinstatement is not a unitary mechanism, but it flexibly adapts to the type of visual content, the dimension of memory being probed, and the availability of linguistic support. Location memory was tied most strongly to trial-specific sensory reinstatement in early visual cortex, whereas color memory recruited an interplay between perceptual and language-related regions, with richer color naming sharpening reinstated representations. Vividness and precision were likewise supported by distinct patterns of reinstatement, reflecting different contributions of sensory and semantic systems. Together, these results underscore how perceptual and linguistic systems dynamically interact to construct detailed and vivid visual memories.

## Acknowledgements

X.W. and R.C. were funded by the Einstein Foundation Berlin (EPP-2017-423, R.C.) and National Institute of Health (NIA 1RF1AG066901, R.C.). We would like to express our deepest gratitude to Paola Gega, Mindia Winkert, Yasamin Azizi and Teuta Dzaferi for their substantial assistance in data collection and Loris Naspi for useful discussions on data processing and analysis. We also acknowledge the use of ChatGPT (OpenAI) for assistance with language refinement.

